# Recombination peaks and fast evolution

**DOI:** 10.1101/2025.03.06.641881

**Authors:** Kristina Crona

## Abstract

Genetic recombination can speed up or slow down evolution toward the global optimum. For analyzing mutational trajectories to the optimum it is conventional to study peaks in the fitness landscapes. However, such mutational peaks cannot provide a complete picture of the interplay between mutations, recombination and selection. Here we use recombination peaks as an additional tool. Informally, a recombination peak is a set of genotypes such that no allele shuffling among members can improve average fitness. If conventional peaks are optimal with respect to mutations, then recombination peaks are optimal with respect to recombination. The approach applies to general fitness landscapes with an arbitrary number of alleles and loci, and our experiments involve up to five alleles.

One of the most frequently observed patterns of epistasis in nature is diminishing returns epistasis, i.e., the combined effect of beneficial mutations is smaller than a multiplicative expectation would predict. Recombination tends to speed up adaptation toward a peak whenever there is a systematic pattern of diminishing returns epistasis near the peak. The opposite is true for increasing returns epistasis. Mixed curvature is more challenging to analyze. Recombination peaks provide a language for describing effects of recombination, and threshold results are discussed accordingly. The experiments suggest that recombination is more likely to speed up adaptation for larger fitness landscapes.

## 1. Introduction

Evolutionary aspects of recombination have been studied for over a hundred years with the goal to understand why recombination is widespread in nature. The famous paradox of recombination refers to that gene shuffling is nearly universal, although nobody has found a universal advantage of recombination. A related and in a sense more practical problem is the effect of recombination. Recombination can speed up or slow down evolution. If a new viral or bacterial variant has a higher recombination rate than the original type, it would be valuable to know whether one should expect faster or slower evolution, or perhaps no change at all. It should be noted that fast evolution does not imply that recombination as a mechanism is selected for. The evolutionary advantage or disadvantage of recombination is usually modeled by recombination modifiers, for more background on recombination see e.g., Otto and Lenormand (2002). Here, the goal is to analyze the effect of recombination, regardless of its evolutionary origin or any other evolutionary aspects of the mechanism as such.

A fundamental observation about recombination is that the mechanism has no effect in the absence of linkage disequilibria i.e., for an already well-mixed population additional gene shuffling has no impact. The terminology can be illustrated for a biallelic two-locus system with genotypes 00, 10, 01, 11. For a population in equilibrium the product of the proportions of the extreme genotypes (00 and 11) equals the product of the proportions of the intermediate genotypes (10 and 01). If the extreme genotypes are underrepresented, then recombination tends to generate the extreme genotypes, and the opposite is true if the extreme genotypes are overrepresented. A common reason for linkage disequilibria is epistasis, or gene interactions. Epistasis measures deviations from multiplicative fitness, where the fitness of a genotype is defined as its expected contribution to the next generation.

For an evolutionary process from the wild-type 00 to the double mutant 11, where each mutation 0 → 1 increases fitness, recombination tends to slow down adaptation for increasing returns epistasis, i.e., if the fitness of 11 exceeds a multiplicative expectation (evolutionary speed is measured as the expected number of generations from the wild-type population 00 until fixation at the optimal genotype 11). That should be intuitively clear from considering the linkage disequilibria caused by the epistasis. By a similar argument, recombination tends to speed up adaptation for diminishing returns epistasis. The impact of recombination is more challenging to describe for several alleles and loci. For an overview of the problem it is natural to consider fitness landscapes. A fitness landscape for an *L*-locus system with *k* alleles at each locus is determined by the fitness values of all *k*^*L*^ genotypes. If the fitness of a genotype is interpreted as a height coordinate, then an evolutionary process for a population can be represented as a walk in the fitness landscape where each step increases the height, i.e., each step represents a single point mutation that increases fitness. A genotype is a peak if it has higher fitness than its mutational neighbors.

For predicting the effect of recombination, one needs to consider accessible walks in the fitness landscape. The rank order of genotypes with respect to fitness determines possible walks. The second key factor is the curvature of the landscape. An obvious complication is that the majority of larger fitness landscapes have mixed curvature with both diminishing and increasing returns epistasis. In addition to rank orders and curvature, population parameters can make a difference. For instance, the effect of recombination is highly sensitive to population size for some landscapes with poor peak accessibility (Crona, 2018).

## 2. Results

Evolutionary theory has historically mostly been focused on the biallelic case. One reason is that many important empirical sets, ranging from bacterial variants associated with antimicrobial drug resistance, to the human genome, are effectively biallelic. However, a complete study of evolutionary potential and constraints has to take into account that the alphabets of life have more than two letters. We start with some foundational observations for the multiallelic case before analyzing recombination.

### 2.1. Foundational multiallelic theory

Throughout the paper, we consider systems with *L* loci and *k* alleles at each locus. Let Σ = {0, 1, …, *k* − 1}. A genotype is an element in Σ^*L*^. A fitness landscape *w* : Σ^*L*^ → ℝ assigns a non-negative fitness value *w*_*g*_ to each genotype *g*. The Hamming distance between two genotypes is defined as the number of loci where they differ. Two genotypes are mutational neighbors if their Hamming distance is 1.

The concept of reciprocal sign epistasis from the biallelic theory can be generalized to the multiallelic case.

#### Definition 2.1.

A fitness landscape has reciprocal sign epistasis if there exists a 2-face whose two genotypes with highest fitness have Hamming distance 2.

The definition is equivalent to the standard notion in the biallelic case. For a multiallelic example, a fitness landscape *w* has reciprocal sign epistasis if both *w*_1245_ and *w*_1236_ exceed the fitness values *w*_1246_ and *w*_1235_ of the intermediates. A classical result in the biallelic theory is that fitness landscapes with multiple peaks have reciprocal sign epistasis (Poelwijk, 2007). This observation can be extended to the multiallelic case.

#### Theorem 2.2.

*A fitness landscapes with muliple peaks has reciprocal sign epistasis. Proof*. Let *w* : Σ^*L*^ → ℝ be a fitness landscape peaks and *p* and *q*, where

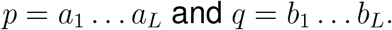

Let

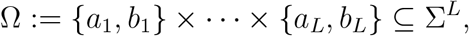

and let 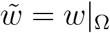 be the restriction of *w* to the domain Ω. Since p and q are global maxima of *w* they remain peaks of 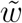. If *a*_*i*_ = *b*_*i*_ for some *i* then the *i*:th coordinate is simply a place holder for *g* ∈ Ω. Consequently 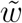 can be considered a fitness landscape for a biallelic *L*′-locus system, where *L*′ is the number of coordinates such that *a*_*i*_ ≠ *b*_*i*_.

Since 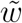 is a biallelic landscape with two peaks it has reciprocal sign epistasis, i.e. there exists a 2-face such that its two genotypes of highest fitness *g*^1^ and *g*^2^ has Hamming distance 2. Since Ω is a face of Σ^*L*^, the genotypes *g*^1^ and *g*^2^ also belong to a 2-face of Σ^*L*^, and their Hamming distance in Σ^*L*^ is likewise 2. By definition, *w* has reciprocal sign epistasis. □

The maximal number of peaks for a biallelic *L*-locus system is 2^*L*−1^, i.e., half of the genotypes can be peaks. This fact was first observed in (Haldane, 1931).

For the multiallelic case we start with a recursive construction of potential peak sets. Such sets cannot include mutational neighbors. For *k* = 2 let

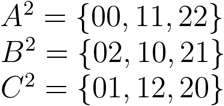

Cne can construct fitness landscapes with exactly three peaks by assigning either *A*^2^, *B*^2^ or *C*^2^ as the set of peaks.

Let

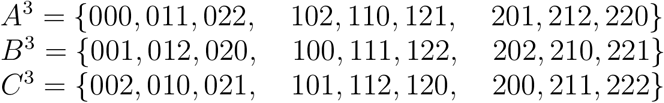

For short, we use the notation:

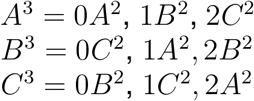

Define *A*^*r*^, *B*^*r*^ and *C*^*r*^ recursively as

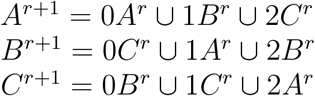

#### Lemma 2.3.

*Any two elements in either A*^*n*^, *B*^*n*^ *or C*^*n*^ *have Hamming distance at least 2, and A*^*n*^ ∩ *B*^*n*^ = *A*^*n*^ ∩ *C*^*n*^ = *B*^*n*^ ∩ *C*^*n*^ = ∅ *for n* ≥ 2.

*Proof*. By inspection, the claim holds for *n* = 2. Assume that the claim holds for *n* = *r*. Then it is immediate that the nine sets

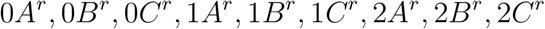

are pairwise disjoint. It follows that *A*^*r*+1^ ∩*B*^*r*+1^ = *A*^*r*+1^ ∩*C*^*r*+1^ = *B*^*r*+1^ ∩*C*^*r*+1^ =∅.

Consider a pair of elements in *A*^*r*+1^ = 0*A*^*r*^ ∪1*B*^*r*^ ∪2*C*^*r*^. If both of them belong to 0*A*^*r*^, then it is clear that they have Hamming distance at least 2. If one of them belongs to 0*A*^*r*^ and the other to 1*B*^*r*^, then they differ in the first coordinate and, since *A*^*r*^ ∩ *B*^*r*^ = ∅, in at least one more coordinate, so that they have Hamming distance at least 2. Analogously, any pair of elements in *A*^*r*+1^ have Hamming distance at least 2, and the same argument holds for *B*^*r*+1^ and *C*^*r*+1^. The result follows by induction. □

A similar construction works in general

For *k* ≥ 1, let

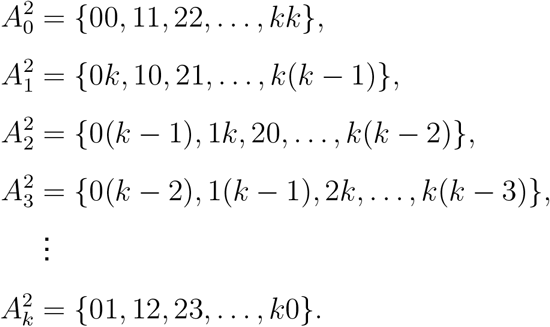

Define 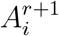 recursively as

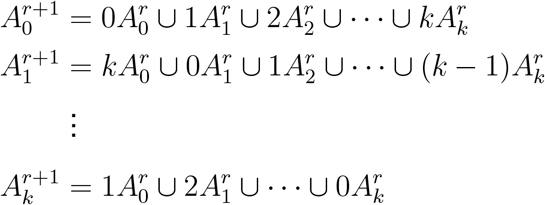

#### Lemma 2.4.

*Let* 0 ≤ *i, j* ≤ *k, where k* ≥ 1. *Any two elements in* 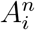 *have Hamming distance at least 2, and the intersection* 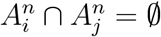 *if i /*= *j for any n* ≥ 2.

*Proof*. The result follows by induction with an argument similar to the previous lemma. □

#### Observation 2.5.

Consider system with *L* locus and *k* alleles at each locus. An upper bound for the number of peaks is *k*^*L*−1^, and the bound is exact.

*Proof*. One can partition the set of peaks into *k* sets *S*_*i*_, where *i* is the first coordinate of the genotype.

If *g* ∈ *S*_*i*_ and *h* ∈ *S*_*j*_, then *g* = *ig*′ and *h* = *jh*′ where *g*′ = *h*′, since otherwise *g* and *h* would be mutational neighbours. The number of elements in an *L* − 1-locus system where each locus has *k* alleles is *k*^*L*−1^. Consequently *k*^*L*−1^ is an upper bound for the number of peaks. The bound is exact, since one can construct a graph such that all elements in 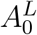 are peaks by directing all edges toward the elements in 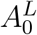. □

#### Remark 2.6.

The set 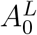 is closely related to classical objects in coding theory. In particular, sets of this type correspond to codes that achieve the maximal size *k*^*L*−1^ under minimum Hamming distance constraints. Under the assumption stated in the observation, an alternative to 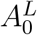 is the set that consists of all sequences that sum up to 0 modulo (*k* − 1). For *L* = 3 and *k* = 2 the set consists of 002, 011, 101, and so forth. This example is standard in coding theory.

### 2.2. Rank orders and Curvature

The biallelic two-locus landscape discussed in the introduction has a simple rank order. As a complement, one can consider the effect of recombination for chaotic rank orders.

#### Example 2.7.

Consider adaptation from 000 to 111 for a finite population with fitness values

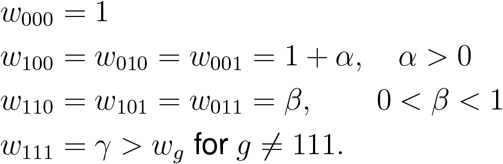

Figure 2.2 shows the fitness graph (Crona et al., 2013; Krug and De Visser, 2009) for the system. Adaptation from 000 to 111 is difficult because of the suboptimal peaks 100, 010, 001, at least if double mutations are rare. Recombination has the potential to speed up adaptation because of *peak jumping*, i.e., that local peaks are replaced by the global peak, depending on *α, β, γ* and population parameters. Not surprisingly, for simulations where *α* is replaced by −*α*, recombination is no longer beneficial. Specifically, that is the case for *α* = *β* = 0.1, *λ* = 3 (Crona, 2018). It is easy to verify that the replacement of *α* by −*α* leaves the curvature unchanged. (See Section 5.1 for a discussion about curvature for biallelic 3-locus systems.)

**FIGURE 1.**
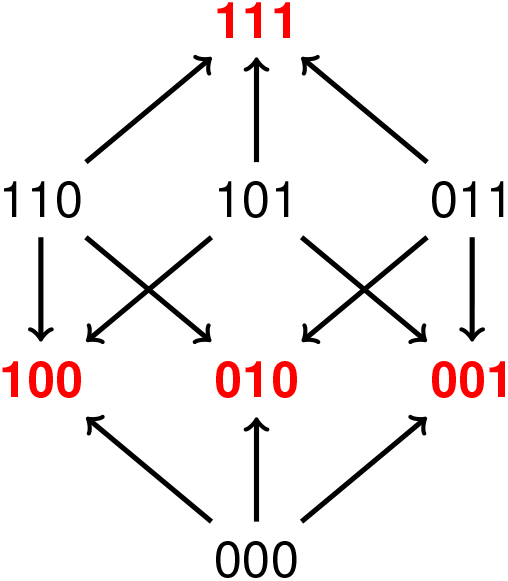
The three suboptimal peaks 100, 010, 001 make all paths from 000 to 111 inaccessible. Depending on population parameters, recombination can be beneficial because of peak jumping, meaning that the suboptimal peaks are replaced by the global peak through repeated allele shuffling.

**FIGURE 2.**
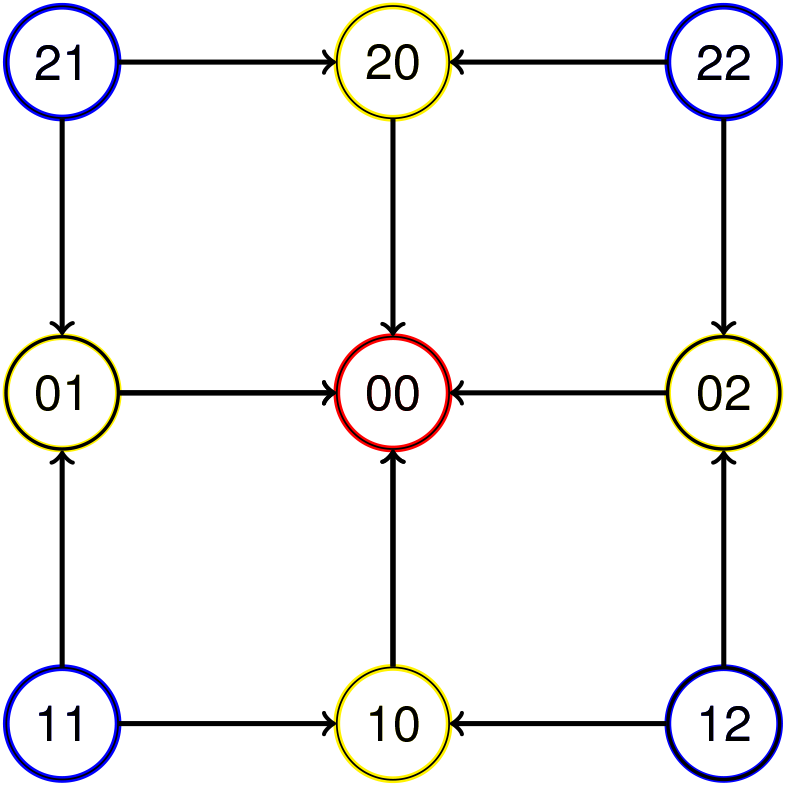
Fitness increases for each mutation 1 ↦ 0 and 2 ↦ 0. Initially the population is distributed on the blue marked genotypes 21, 22, 11, 12, and eventually the population goes to fixation at the peak 00. The genotype 00 is the global peak. Each arrow points toward the genotype with higher fitness. For clarity, arrows are included for mutational neighbors with exception for the ones that have the same distance to the global peak (for instance, no arrow connects 10 and 20).

The example shows that the effect of recombination cannot be predicted from curvature type alone. As clear from the introduction, neither can the effect be determined from rank orders of genotypes with respect to fitness. A complete analysis would need to capture the interplay between rank order aspects and curvature.

In order to isolate the effect of curvature it is useful to consider rank orders that are as simple as possible. To avoid some subtle aspects due to stochasticity (Barton and Otto, 2005; Hill and Robertson, 1966), populations are also assumed to be infinite and well mixed throughout the remainder of the paper. The biallelic two-locus system discussed in the introduction, with genotypes 00, 10, 01, 11 and where each mutation 0 ↦ 1 increases fitness, is an important reference for larger fitness landscapes with simple rank orders. In more detail, let *f*_*g*_ denote fitness and *w*_*g*_ log fitness for a genotype *g*.

Then

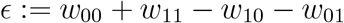

is zero if fitness is multiplicative, the system has diminishing returns epistasis if *ϵ* < 0, and increasing returns epistasis if *ϵ* > 0. Under the stated assumptions recombination speeds up adaptation from 00 to 11 for *ϵ* < 0 and slows down adaptation for *ϵ* > 0 whereas recombination has no effect if *ϵ* = 0 (as explained in the introduction). The (Hamming) distance between two genotypes is defined as the number of loci where the genotypes differ, and two genotypes are called mutational neighbors if their distance is one.

In general, the rank orders are assumed to satisfy that the landscapes are single peaked and that each step toward the global peak increases fitness. Since larger fitness landscapes tend to have mixed curvature, i.e., both diminishing returns and increasing returns epistasis, it is more challenging to analyze the effect of recombination. Figure 2.2 shows a fitness graph for a two-locus system with three alleles. Each arrow in the figure points toward a genotype of higher fitness. The peak 00 (marked red) has mutational neighbors 10, 20, 01, 02 (marked yellow), and the genotypes on distance two (marked blue) are 11, 12, 21, 22. By assumption, there are no obstacles in the landscape that prevent walks from an arbitrary genotype to the peak 00. For a clean presentation only arrows between genotypes of different distances to the peak are included in the figure, whereas a complete fitness graph has arrows between all pairs of mutational neighbors.

For the class of fitness landscapes described by the figure, assume that the initial population is equally distributed on the genotypes of distance two from the peaks, and for simplicity that such genotypes have fitness 1. Extrapolating from the biallelic case, one expects that recombination speeds up adaptation for low values of *f*_00_ and slows it down for high values of *f*_00_. If all intermediates have exactly the same fitness (Table 1), then the effect of recombination is indeed very similar to the biallelic two-locus case. However, if the intermediates have different fitness values, then the curvature is mixed for some values of *f*_00_. For the experiments (Table 3) systematic diminishing returns epistasis implies a speed advantage for recombination and increasing returns epistasis a disadvantage, which should not be surprising. The more remarkable finding is that the threshold is relatively high within the interval for mixed curvature, specifically about 3/4 from the left end of the mixed curvature interval (similar results can be found in Tables 4-10). An intuitive explanation for the relatively high thresholds is that evolution tends to use high fitness intermediates rather than low fitness intermediates. The finding suggests that recombination is more likely to speed up adaptation for larger fitness landscapes as compared to the biallelic two-locus case. The threshold problem discussed for two loci can be extended to general landscapes with simple rank orders.

**TABLE 1.** Initially the population consists of 00 genotypes, and eventually the population goes to fixation at 11 after the numbers of generation shown with and without recombination (bottom line). Recombination speeds up adaptation for diminishing returns epistasis (left) and slows it down for increasing returns epistasis (middle), whereas there is no effect if *ϵ* = 0.

| $g$ | $f_g$ | $g$ | $f_g$ | $g$ | $f_g$ |
| --- | --- | --- | --- | --- | --- |
| 00 | 1 | 00 | 1 | 00 | 1 |
| 10 | 1.1 | 10 | 1.1 | 10 | 1.1 |
| 01 | 1.1 | 01 | 1.1 | 01 | 1.1 |
| 11 | 1.14 | 11 | 1.3 | 11 | 1.21 |
| Epistasis | $\epsilon < 0$ | Epistasis | $\epsilon > 0$ | Epistasis | $\epsilon = 0$ |
| with rec. | 391 | with rec. | 236 | with rec. | 274 |
| no rec. | 690 | no rec. | 172 | no rec. | 274 |

**TABLE 2.** The experiment is similar to Table 1 except that there are three alleles 0, 1, 2. Here 00 is the global optimum and the starting point is a population equally distributed on 11, 12, 21, 22. The results differ only marginally from Table 1.

| $g$ | $w_g$ | $g$ | $f_g$ | $g$ | $f_g$ |
| --- | --- | --- | --- | --- | --- |
| 00 | 1.14 | 00 | 1.3 | 00 | 1.21 |
| 10 | 1.1 | 10 | 1.1 | 10 | 1.1 |
| 01 | 1.1 | 01 | 1.1 | 01 | 1.1 |
| 20 | 1.1 | 20 | 1.1 | 20 | 1.1 |
| 02 | 1.1 | 02 | 1.1 | 02 | 1.1 |
| 11 | 1 | 11 | 1 | 11 | 1 |
| 12 | 1 | 12 | 1 | 12 | 1 |
| 21 | 1 | 21 | 1 | 21 | 1 |
| 22 | 1 | 22 | 1 | 22 | 1 |
| Epistasis | $\epsilon < 0$ | Epistasis | $\epsilon > 0$ | Epistasis | $\epsilon = 0$ |
| with rec. | 399 | with rec. | 243 | with rec. | 281 |
| no rec. | 710 | no rec. | 177 | no rec. | 281 |

**TABLE 3.** The table shows the fitness values for the genotypes in a twolocus system with three alleles. By assumption, the original population is distributed equally on the four genotypes with maximal distance from the peak 00. Experiments show that the recombination threshold for, i.e., the value such that evolution is equally fast with and without recombination, is *f*_00_ = 1.38. Systematic diminishing returns epistasis requires that *f*_00_ < 1.1^2^ = 1.21, and systematic increasing returns epistasis that *f*_00_ > 1.2^2^ = 1.44.

| $g$ | $f_g$ |
| --- | --- |
| 10 | 1.1 |
| 01 | 1.1 |
| 20 | 1.2 |
| 02 | 1.2 |
| 11 | 1 |
| 12 | 1 |
| 21 | 1 |
| 22 | 1 |
| diminishing returns | $f_{00} < 1.21$ |
| increasing returns | $f_{00} > 1.44$ |
| $r$-threshold | $f_{00} = 1.38$ |

**TABLE 4.** The table shows the fitness values for the genotypes in a twolocus system with five alleles. By assumption, the original population is distributed equally on the 16 genotypes with maximal distance from the peak 00. Experiments show that the recombination threshold for, i.e., the value such that evolution is equally fast with and without recombination, is *f*_00_ = 2.6. From the theory, the threshold is between 1.1 *×* 1.2 = 1.32 and 1.7 *×* 1.8 = 3.06, so that the threshold falls closer to the upper bound. For a reference, the condition that all sets {00, *g* : *g ∈ G*_2_} are recombination peaks implies that *f*_00_ ≥ 3.06.

| $g$ | $f_g$ |
| --- | --- |
| 10 | 1.1 |
| 01 | 1.2 |
| 20 | 1.3 |
| 02 | 1.4 |
| 30 | 1.5 |
| 03 | 1.6 |
| 40 | 1.7 |
| 04 | 1.8 |
| $g \in G_2$ | 1 |
| $r$-threshold | $f_{00} = 2.6$ |

#### Definition 2.8.

*The threshold problem* is defined for a single peaked fitness landscape that satisfy the following conditions:

i. The initial population consists of genotypes of distance *L* from the peak *p*, and the population is uniformly distributed over such genotypes.
ii. (a) If there are more than two alleles at each locus, then each mutation *a* ↦ 0 for *a ≠* 0 increases fitness, (b) for biallelic systems, each mutation 0 ↦ 1 increases fitness.
iii. Unless otherwise stated, we also assume *f*_*g*_ = 1 for all *g* of distance *L* from the peak.

Under the stated assumptions, the goal is to find the threshold *T* such that recombination speeds up adaptation if *f*_*p*_ < *T* and slows it down if *f*_*p*_ *> T* (if such a threshold exists).

The definition is intended to be relevant for the end behavior of adaptation. A rugged landscape can still be smooth near a peak, and then the conditions (i) and (ii) apply for the last few steps of the adaptation. The somewhat inconsistent notation in condition (ii) depends on that it is convenient to use zeros for the global peak in the multiallelic case, whereas the convention in the biallelic case is that zeros denote the wild-type alleles. Note that Condition (iii) is not of any theoretical importance, but included for easy comparisons of experiments. Throughout the remainder of the paper the landscapes are assumed to satisfy Conditions (i)-(iii), unless otherwise stated. The main purpose with the experiments was to better understand recombination thresholds. For additional details about the the assumptions for the experiments, see Section 5.2

### 2.3 Recombination peaks for genotypes of distance two

Landscapes that satisfy the conditions in Definition 2.8, have either systematic diminishing returns epistasis, systematic increasing returns epistasis, or mixed curvature (as illustrated for landscapes with two loci and three alleles). For a more detailed analysis it is convenient to consider sets of genotypes that are optimal with respect to recombination. We call such sets *recombination peaks*. The word “peak” merely refers to optimality (no mountain top metaphor is intended).

For minimal landscapes, the set {10, 01} is a recombination peak exactly if *w*_11_ +*w*_00_ < *w*_10_ + *w*_01_. That should make sense since allele shuffling 10 + 01 ↦ 11 + 00 decreases mean fitness. Similarly, increasing returns epistasis is equivalent to that {00, 11} is a recombination peak.

Analogously, for biallelic 3-locus landscapes (systematic) diminishing returns epistasis is equivalent to that

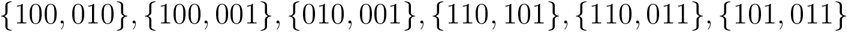

are recombination peaks (notice that the genotypes in each pair have distance two, and that they have same distance to the peak 111), and increasing returns epistasis to that

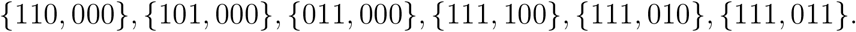

are recombination peaks. For two locus and three alleles, diminishing returns epistasis means that {10, 01}, {10, 02}, {01, 20}, {02, 20}are recombination peaks, and increasing returns epistasis that {00, 11}, {12, 01}, {21, 00}, {22, 00}are recombination peaks. Diminishing and increasing returns epistasis can be expressed similarly for any number of loci and alleles. Note also that a pair of genotypes with distance two constitutes a recombination peak exactly if exchanging alleles at the two loci where they differ does not result in higher mean fitness (a general definition of recombination peaks will be discussed in Section 2.4).

The following claim appears obvious. However, we do not have a theoretical proof.

#### Conjecture 1

*Assume that an L-locus fitness landscape with peak p and with an arbitrary number of alleles satisfies the conditions in Definition 2.8*.

*Consider the pairs of genotypes such that*

1. *the distance to p is the same for both genotypes within the pair, and it is less than L*,
2. *the distance between the genotypes in the pair is two*.

*If all such pairs are recombination peaks, then recombination speeds up adaptation. If none of the pairs are recombination peaks, then recombination slows down adaptation*.

For biallelic landscapes, the condition that all the pairs which satisfy (1) and (2) are recombination peaks is equivalent to that the landscape has universal negative epistasis (as defined in Krug and Oros (2024)), and the opposite extreme to that the landscape has universal positive epistasis (Crona et al., 2023).

##### Example 2.9.

Assume that the *f*_11_ = *f*_12_ = *f*_21_ = *f*_22_ = 1, and that the intermediates have fitness

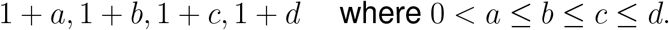

By assumption, *f*_00_ > 1 + *d*. Let

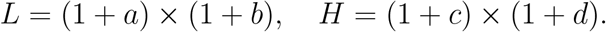

Then all pairs {10, 01}, {10, 02}, {01, 20}, {02, 20} are recombination peaks if *f*_00_ < *L* and that none of them if *f*_00_ *> H*. According to the conjecture, recombination speeds up adaptation if *f*_00_ < *L* and slows it down if *f*_00_ *> H*. However, the conjecture does not apply if *L* < *f*_00_ < *H*. For an analysis of the exact recombination threshold it seems natural to consider some notion of average curvature. A simple measure [which agrees with the threshold in the case *a* = *b* = *c* = *d*] is

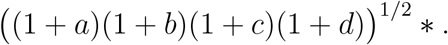

Experiments show that the threshold depends on how the genotypes are positioned in the fitness graph (Table 5). Specifically, the threshold is greater for

**TABLE 5.** The fitness values for Tables a) and b) are the same, except for a permutation of fitness values for the intermediate genotypes, and similarly for Tables c) and d). The tables show that the permutations have a substantial impact on the recombination thresholds.

|  |  |  |  |  |  |
| --- | --- | --- | --- | --- | --- |
| a) | $g$ | $f_g$ | b) | $g$ | $f_g$ |
|  | 10 | 1.4 |  | 10 | 1.4 |
|  | 01 | 1.4 |  | 01 | 1.8 |
|  | 20 | 1.8 |  | 20 | 1.4 |
|  | 02 | 1.8 |  | 02 | 1.8 |
|  | 11 | 1 |  | 11 | 1 |
|  | 12 | 1 |  | 12 | 1 |
|  | 21 | 1 |  | 21 | 1 |
|  | 22 | 1 |  | 22 | 1 |
| $r$-threshold | | $f_{00} = 2.93$ | $r$-threshold | | $f_{00} = 2.52$ |
| c) | $g$ | $f_g$ | d) | $g$ | $f_g$ |
|  | 10 | 1.3 |  | 10 | 1.3 |
|  | 01 | 1.4 |  | 01 | 1.6 |
|  | 20 | 1.5 |  | 20 | 1.4 |
|  | 02 | 1.6 |  | 02 | 1.5 |
|  | 11 | 1 |  | 11 | 1 |
|  | 12 | 1 |  | 12 | 1 |
|  | 21 | 1 |  | 21 | 1 |
|  | 22 | 1 |  | 22 | 1 |
| $r$-threshold | | $f_{00} = 2.32$ | $r$-threshold | | $f_{00} = 2.16$ |

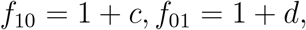

as compared to

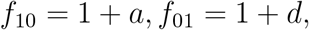

provided *a* < *c* < *d*. Intuitively, if the highest fitness values are associated with a pair of genotypes that can produce the global peak through recombination, then recombination is more likely to speed up adaptation.

For instance, with *a* = 0.3, *b* = 0.4, *c* = 0.5 and *d* = 0.6, the recombination threshold is 2.32 or 2.16, depending on how the intermediates are placed in the fitness graph. For comparison, the lower bound from the conjectures is *L* = 1.82 and *H* = 2.4, and the measure * equals 2.08. The experiments illustrate that the thresholds (2.32 or 2.16) in both experiments are relatively high in comparison with *H, L* and *, and secondly that the positions of the intermediates in the fitness graph have an impact.

The expression * can be generalized to an arbitrary fitness landscape that satisfies Definition 2.8, specifically

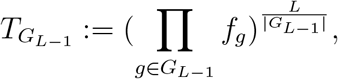

where *G*_*k*_ denote the genotypes of distance *k*. One can perhaps consider 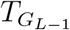 as the best estimate if the only available information is fitness values of genotypes in *G*_*L*−1_ and *G*_*L*_. For a biallelic landscape with multiplicative fitness for all genotypes except the peak, the recombination threshold is exactly 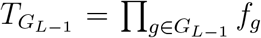. However, the estimate is a poor predictor unless fitness is nearly multiplicative.

##### Example 2.10.

Under the assumption that

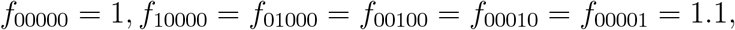

and that the remaining intermediates follow multiplicative expectations except for that

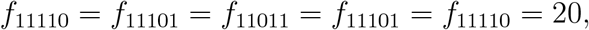

the recombination threshold is *f*_11111_ = 25.8. In other words, recombination speeds up adaptation if 20 < *f*_11111_ < 25.8, and slows down adaptation if *f*_11111_ > 25.8. For comparison 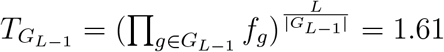.

An estimate based on a comparison between *f*_*p*_ and the wild-type versus *all intermediates* has a better chance to be relevant for the recombination threshold. To that end it makes sense to consider recombination peaks for an arbitrary set of genotypes.

### 2.4. General recombination peaks

A recombination peak is a set of genotypes that is optimal with respect to recombination. One can verify that the set {111, 000} is as a recombination peak if

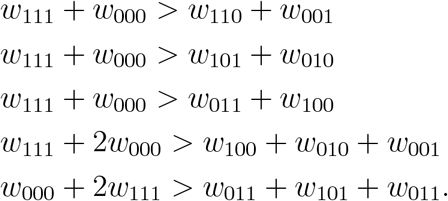

It is immediate that the three first inequalities are necessary. The bottom inequality is also necessary since one example of (repeated) allele shuffling starting out with the three genotypes 000, 111, 111 is

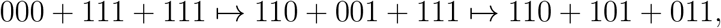

and the remaining inequality can be explained similarly. More background on how to determine if a set constitutes a recombination peak is provided Section 2.6.

For fitness landscapes that satisfy i-iii) such that {111, 000} is a recombination peak, it is fair to say that the peak genotype 111 has high fitness as compared to all the intermediates. Our experiments supports that recombination slows down adaptation whenever {111, 000} is a recombination peak (Tables 6-9).

**TABLE 6.** The fitness values are fixed for 000, 100, 010, 001, 111 (Table 6a). The fitness values for the three double mutants are the same in each experiment. The key assumption is that the fitness *f*_111_ equals the multiplicative expectation based on the fitness values *f*_000_, *f*_100_, *f*_101_, *f*_001_. Table 6b) shows how the effect of recombination varies depending on the fitness values for the three double mutants. The number of generations until fixation with and without recombination is given. The table also indicates whether or not {000, 111} is a recombination peak. Notice that fitness is multiplicative exactly if the double mutations have fitness 1.21.

|  |  |  |  |  |  |  |  |
| --- | --- | --- | --- | --- | --- | --- | --- |
| a) | $g$ | $f_g$ | b) | $f_{110}, f_{101}, f_{011}$ | recomb | no recomb | $\{000, 111\}$ |
| | 000 | 1 | | 1.15 | 261 | 238 | $r$-peak |
| | 100 | 1.1 | | 1.20 | 273 | 258 | $r$-peak |
| | 010 | 1.1 | | 1.21 | 278 | 278 | $1.21 = 1.1^2$ |
| | 001 | 1.1 | | 1.25 | 306 | 407 | not $r$-peak |
| | 111 | $1.331 = 1.1^3$ | | 1.30 | 435 | 1024 | not $r$-peak |

One can define recombination peaks for an arbitrary number of alleles and loci. We define the allele counter *C*^*j*^(Σ*w*_*g*_) as the vector that counts how many times the allele *j* occurs in total.

1. *C*^*j*^(*w*_*g*_) is the vector that shows how many time allele *j* occurs (0 or 1) at each locus of *g*,
2. *C*^*j*^(Σ*w*_*g*_) = Σ*C*^*j*^(*w*_*g*_).

For instance, *C*^0^(*w*_10_) = [0, 1] and *C*^0^(*w*_01_) = [1, 0], and from Condition 2,

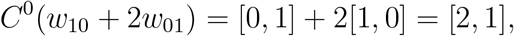

and similarly *C*^1^(*w*_10_ + 2*w*_01_) = [1, 2] and *C*^2^(*w*_10_ + 2*w*_01_) = [0, 0]. For determining if a set of genotypes is a recombination peak one needs to compare sets of genotypes that have the same allele counts.

#### Definition 2.11.

A set *S ⊂* {0, …, *k*}^*L*^ is a recombination peak if for any genotypes *g*_1_, …, *g*_*m*_ *∈ S* and 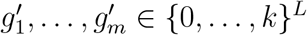 the following condition holds:

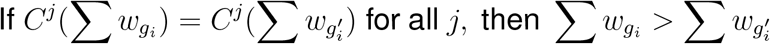

In other words, the sum of the fitnesses for genotypes from a recombination peak exceeds the sum from genotypes with the same allele count. Readers familiar with Beerenwinkel et al. (2007) should notice that recombination peaks are closely related to authors’ concept of a fittest populations (however, note that recombination peaks are *sets* of genotypes).

The following result is immediate:

#### Observation 2.12.

If the set of all genotypes from a population constitutes a recombination peak, then the population cannot increase its average fitness through a sequence of recombination events.

The observation is more or less a tautology but it is less obvious how to determine whether or not a set of genotypes is a recombination peak. For solving the problem one can apply theory on triangulations of polytopes (see Section 2.6).

The following claim, stated as an open problem has some support from experiments. If correct, the claim provides an upper bound for the recombination threshold.

### 2.5. Open problem

*Proposed upper bound for the recombination threshold: For a fitness landscape that satisfies the conditions in Definition 2.8 let p denote the global peak of the fitness landscape. If for all g* ∈ *G*_*L*_ *the pair* {*p, g*} *is a recombination peak, then recombination slows down adaptation*.

The condition stated in the proposed bound simply means that *f*_*p*_ must be sufficiently high as compared to intermediates. For the threshold experiments on mixed curvature (Tables 3-10) the thresholds were close to the proposed upper bound in several cases, and sometimes considerably lower. The threshold problem, at least in the form stated here, is primarily interesting for relatively few loci (otherwise the assumptions would seem less natural). If the proposed bound is not correct, it may still work as a rule of thumb for relatively small fitness landscapes.

Theoretical work on recombination for several loci has primarily considered selection of recombination modifiers (Barton, 1995), which is a problem different from the effect of recombination (Otto and Lenormand, 2002). Some of the fundamental insights from classical work could still be relevant for theoretical questions on recombination thresholds.

### 2.6. Recombination and triangulations of polytopes

The following observation is one of the most precise results on recombination for a general fitness landscapes. The observations rephrases theory developed in (Beerenwinkel et al., 2007). In brief, one can determine whether or not a population is “perfectly combined” by using theory on triangulations (De Loera et al., 2010)..

#### Observation 2.13.

Let *w* be a fitness landscape.

i. One can determine if a population is a recombination peak by checking the signs of a set of forms in the variables *w*_*g*_. In particular, for a biallelic 2-locus system one needs to check the sign of the form *ϵ* = *w*_00_ + *w*_11_ − *w*_10_ − *w*_01_. The forms are known as circuits (see the appendix for a more detailed discussion).
ii. A population is a recombination peak exactly if its genotypes belong to the same simplex in the triangulation induced by the fitness landscape *w*, in the sense described in Beerenwinkel et al. (2007) (see also below for a brief description).

For a biallelic two-locus population one can describe the result in a simple way. Observation (i) translates to that the form

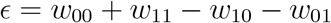

determines if a population is a recombination peak; specifically *ϵ >* 0 implies that {11, 10, 00} and {11, 01, 00} are recombination peaks, whereas *ϵ* < 0 implies that {11, 10, 01} and {10, 01, 00}, are recombination peaks. Observation (ii) translates to that the square with vertices labelled by the genotypes 00, 10, 01, 11 is subdivided to the triangles with vertices 11, 10, 00 and 11, 01, 00 if *ϵ >* 0, and with vertices 11, 10, 00 and 11, 01, 00 if *ϵ* < 0. A population being a recombination peak is equivalent to all genotypes belonging to the same triangle in the induced triangulation.

More generally, for a biallelic *L*-locus system, the genotypes can be considered vertices in an *L*-cube. If *w*_*g*_ is the height coordinate for the genotype *g* and the polytope *P* the convex hull of the resulting point configuration, then the projection of the upper faces of *P* onto the *L*-cube describes the triangulation induced by *w*. As in the two-locus case, the population is a recombination peak if its genotypes belong to the same simplex (the same tetrahedron for *L* = 3, and so forth). Biallelic *L*-locus systems are also discussed in some detail in Crona et al. (2023); Crona (2014).

Here we also consider also *k >* 2 alleles. In brief, let Δ_*k*_ be the *k* − 1 dimensional simplex

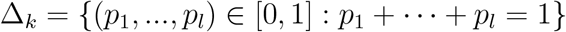

and let 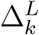 be the *L*-fold direct product of simplices Δ_*k*_. Then the genotypes can be identified with the vertices of the polytope 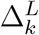. Similar to the biallelic case, *w* induces a triangulation of the 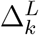.

Consider systematic increasing returns epistasis for biallelic *L*-locus system in our setting. The condition implies that the fitness landscape induces a specific triangulation known as the stair-case triangulation (Crona et al., 2023) For *L* = 3, the triangulation is

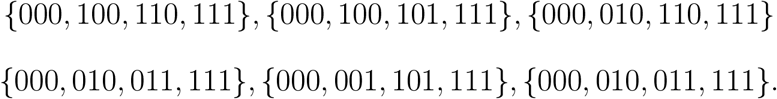

One can ask if a similar result holds for *k >* 2 alleles, Below is a counterexample.

#### Example 2.14.

The following fitness landscapes have systematic increasing returns epistasis.

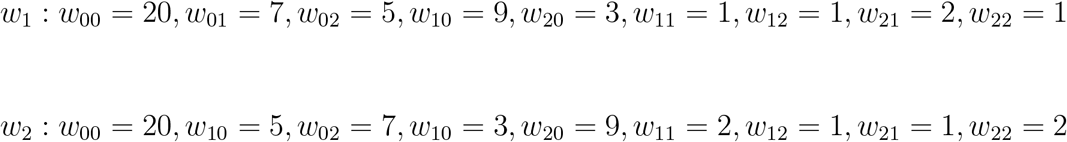

The triangulation for *w*_1_ is

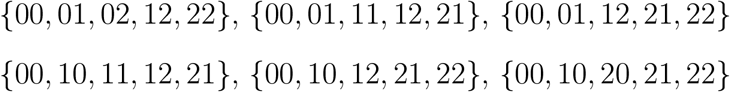

The triangulation for *w*_2_ is

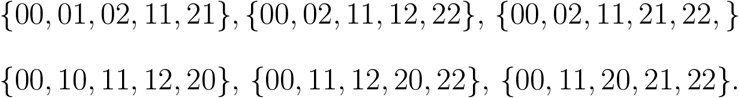

The two triangulations are of different combinatorial types (as explained below), which shows that the condition of increasing returns epistasis does not determine the triangulation type in general.

For analyzing the triangulations induced by *w*_1_ and *w*_2_ in the example, one can use theory that relates triangulations of Δ^2^ × Δ^2^ and lozenge tilings of triangles, see Chapter 9.2 in De Loera et al. (2010). The connection to tilings is known as the Cayley-trick. Figure 9.57 in Chapter 9 shows all combinatorial types of triangulations of Δ^2^ *×* Δ^2^ as lozenge tilings of a triangle. The triangulation of the fitness landscape *w*_1_ from the example corresponds to the second type from the top in the figure, and the triangulation of *w*_2_ to the fourth type from the top.

## 3 Discussion

Recombination can speed up or slow down adaptation. Our experiments suggest that recombination speeds up adaptation for many larger fitness landscapes. Several studies have shown that the impact of recombination is difficult to predict from conventional statistics on pairwise epistasis and similar summary statistics (Kouyos et al., 2006). We propose new tools for relating the curvature of the fitness landscape to the impact of recombination. We have demonstrated that neither rank orders alone, nor curvature alone, can predict the impact. For many chaotic rank orders, recombination is not very sensitive for curvature, whereas curvature is the determining factor for simple rank orders. In addition to rank orders and curvature, the impact of recombination also depends on population parameters.

In order to isolate the effect of curvature, our experiments use a rank order that is as simple as possible, i.e., fitness increases for each step toward a global peak, and infinite population size is assumed. In contrast to most studies, the experiments include landscapes with several alleles and loci and allow for initial populations with several genotypes. Already for two-locus landscapes the effect of several alleles is that recombination is more likely to speed up adaptation. One reason is that evolution tends to use high fitness intermediates rather than low fitness intermediates if there are several alternatives. The finding is yet another example of the recurrent observation in evolutionary biology that the walks chosen by adapting populations differ from arbitrary walks in the fitness landscape (Greene and Crona, 2014; Moradigaravand et al., 2014; Draghi and Plotkin, 2013).

The impact of recombination is clear-cut for minimal landscapes in the setting, i.e., for landscapes with four genotypes 00, 10, 01, 11 such that each mutations 0 ↦ 1 increases fitness. Recombination slows down adaptation if the landscape has increasing returns epistasis (that is if the fitness of 11 exceeds a multiplicative expectation) and speeds it up for diminishing returns epistasis. Our experiments support that the analogous result holds for larger landscapes if the curvature is consistent (either diminishing or increasing returns epistasis). However, for most larger landscapes the curvature is mixed which makes the effect of recombination more challenging to predict. A striking observation from experiments is that the outcome does not depend on the average curvature (of any sort). The speed advantage appears to discount low fitness intermediates if there are high fitness alternatives. Another argument in favor of that recombination tends to speed up adaptation is that diminishing returns epistasis have been observed in many empirical studies (Stround and Ratcliff, 2025; Wei and Zhang, 2019; Otwinowski et al., 2018; Lenski, 2017; Kryazhimskiy et al., 2014).

For more quantitative precision, one needs some notion of curvature and to that end we introduced recombination peaks. A recombination peak is a set of genotypes such that no allele shuffling among members of the set can improve average fitness. For instance, for minimal landscapes diminishing returns epistasis means that {10, 01} is a recombination peak and increasing returns epistasis that {11, 00} is a recombination peak. If one keeps the fitness values of all genotypes fixed, except for the peak, then one can ask how the impact of recombination will change by peak fitness. The threshold experiments assume that the initial population is equally distributed on the same distance from the peak (for more details see Section 5.2). For all landscapes, the experiments identified a threshold such that recombination slows down adaptation if the peak fitness exceeds the threshold and otherwise speeds it up. We found that if {*g, p*} is a recombination peak for all *g* on maximal distance from the peak *p*, then recombination slows down adaption. The condition, if true in general, would provide an upper bound for the recombination threshold. Not surprisingly, the threshold was considerably lower for some landscapes.

It would be interesting to have a more precise understanding of recombination thresholds. One open problem is if recombination thresholds exist for all landscapes in the setting. Perhaps more empirically important, the implications of the observed relatively high recombination thresholds for larger fitness landscapes would be interesting to quantify. For most fitness landscapes it seems nearly impossible to predict the recombination thresholds. However, we conjecture that recombination speeds up adaptation for systematic diminishing returns epistasis and slows it down for increasing returns epistasis, regardless of the number of loci and alleles.

## 4 Acknowledgements

Devin Greene produced all code for the evolutionary experiments and for determining triangulations of polytopes.

## 5. APPENDIX

### 5.1. Circuits and triangulations

A general approach to gene interactions based on polytope theory was introduced in Beerenwinkel et al. (2007). For mathematical background, see De Loera et al. (2010).

The approach uses 20 forms in the variables *w*_*g*_ for analyzing the gene interactions for biallelic 3-locus systems, analogous to the sole form *ϵ* for a biallelic two-locus landscape discussed in the main text. For instance, the form

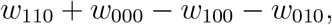

(with last coordinate zero for all genotypes) is relevant for an early phase of the adaptation, and the form

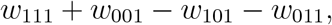

with last coordinate 1 for all genotypes) is relevant for a late phase of adaptation. The 20 forms fall into three categories. Six forms, including

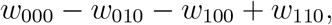

represent faces of the cube; six forms, including

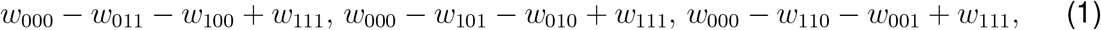

represent squares that cut the cube into two triangular prisms, and eight forms, including

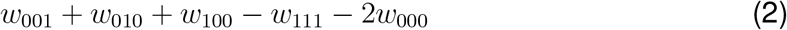

represent bipyramides. For an explicit list of the 20 forms see Beerenwinkel et al. (2007, Page 1352). If the sign of each form (positive or negative) is the same for two biallelic three-locus landscapes, then the gene interactions are considered similar.

Out of the 20 forms exactly five have support in both *w*_000_ and *w*_111_, specifically the five listed in (1) and (2) above. Consequently, one needs to check five inequalities for determining if {000, 111} constitutes a recombination peak, exactly as stated in Section 2.4. The 20 forms constitute the complete set of circuits for a biallelic three-locus system. For a general definition of circuits, let *C* denote the allele counter defined in the main text, and let *C*^*j*^(Σ*a*_*i*_*w*_*g*_) = Σ*a*_*i*_*C*(*w*_*g*_). For *ϵ* = *w*_00_ + *w*_11_ − *w*_10_ − *w*_01_ one gets *C*^0^(*ϵ*) = [1, 1] + [0, 0] − [0, 1] − [1, 0] = [0, 0], *C*^1^(*ϵ*) = [0, 0] + [1, 1] − [1, 0] − [0, 1] = [0, 0].

A circuit can be defined as a linear form in the variables *w*_*g*_ such that

a. the allele counter *C*^*j*^ equals the zero-vector for all *j*,
b. its support (the *w*_*g*_ which appear with non-zero coefficients) is non-empty but minimal with respect to inclusion.

Recent applications of the described approach to interactions in evolutionary biology and ecology includes (Brown et al., 2023; Crona et al., 2023; Eble et al., 2023; Crona, 2020; Eble et al., 2019). A different approach to gene interactions for biallelic systems relies on Walsh coefficients (Weinreich et al., 2013), for mathematical background see also (Greene, 2023), with recent extensions to the multiallelic case (Faure et al., 2024; Greene, 2023; Crona and Greene, 2024).

### 5.2 Experiments

All experiments concern infinite populations that are well mixed. We use a Fisher-Wright type of model where relative offspring proportions are determined by fitness. Mutation is followed by recombination in the reproductive step. Finally the offspring are chosen to be the parents of the subsequent generation. Recombination is modeled so that for the resulting genotype each locus is equally likely to agree with either parent’s allele. The experiment halts if the proportion of the peak genotype exceeds 99.99 percent of the population. The mutation rate is 10^−9^ (however, most results are not sensitive for the mutation rate). The speed of adaptation is measured as the number of generation from wild-type population to fixation at the global peak.

Recall that *G*_*L*_ denotes genotypes of distance *L* from the the peak *p*. The assumptions are the same as the conditions (i)-(iii) stated in Definition 2.8, i.e.,

i. The initial population consists of genotypes of distance *L* from the peak *p*, and the population is uniformly distributed over such genotypes.
ii. (a) If there are more than two alleles at each locus, then each mutation *a* ↦ 0 for *a* ≠ 0 increases fitness, (b) for biallelic systems, each mutation 0 ↦ 1 increases fitness.
iii. Unless otherwise stated, we also assume *f*_*g*_ = 1 for all *g* of distance *L* from the peak.

The experiments in Tables 3-5 and 7-10 study the threshold *T* such that recombination slows down recombination if *f*_*p*_ *> T* and speeds up recombination if *f*_*p*_ < *T*. The main purpose with the experiments was to gain some understanding of the recombination threshold (*r*-threshold).

For Table 3 the landscape has two loci and three alleles. Two intermediates have fitness 1.1 and the other two 1.2. It follows that *f*_00_ < 1.21 implies systematic diminishing returns epistasis and *f*_00_ > 1.44 increasing returns epistasis. Extrapolating from the biallelic two-locus case, one expects that recombination speeds up adaptation if *f*_00_ < 1.21 and slows down adaptation if *f*_00_ < 1.44, but the outcome is less obvious for values inbetween. The recombination threshold is 1.38, clearly much closer to 1.44 (difference - 0.6) than to 1.21 (difference +0.17). As explained in the main text, an intuitive explanation for the relatively high recombination threshold is that evolution tends to use high fitness intermediates rather than low fitness intermediates.

For Table 4 the landscape has two loci and five alleles. By the general assumptions, the original population is distributed equally on the 16 genotypes with maximal distance from the peak 00, all of them having fitness 1. All the intermediates have different fitness values, the two lowest 1.1 and 1.2, and the highest 1.7 and 1.8. It follows that the recombination threshold is expected to fall between 1.1 · 1.2 = 1.32 and1.7· 1.8 = 3.06. The recombination threshold is 2.6, again relatively high.

It should be noted that the outcome is sensitive for details. Table 5 investigates the effect of a permutation of fitness values for the intermediates. In brief, the table shows that if the peak can be produced directly from high fitness intermediates, then the recombination threshold is higher.

Tables 6-10 concerns biallelic three-locus systems. For table 6 the single mutants *f*_100_ = *f*_010_ = *f*_00_ = 1.1, and the peak *f*_111_ = 1.331 are all fixed and agrees with a multiplicative expectation. The three double mutants have the same fitness, and it varies between 1.15 and 1.31 in the experiments. Consequently, the fitness landscape has mixed curvature for all fitness values of double mutants (except if the value is exactly 1.21). The table shows that recombination speeds up adaptation if {000, 111} is a recombination peak (which implies that the double mutants have fitness below the multiplicative expectation), and slows down adaptation otherwise. Recombination speeds up adaptation considerably toward the right end of the interval (435 generations with recombination, versus 1024 generations without recombination).

Table 7 works with assumptions similar to Table 6, and computes recombination thresholds. Tables 7-10 support the claim that recombination slows down adaptation if {000, 111} is a recombination peak, and shows that the threshold is considerably lower under some circumstances. Table 10 shows that recombination can speed up adaptation for landscapes with substantial increasing returns epistasis, as long as the very last step of adaptation is almost flat.

**TABLE 7.**
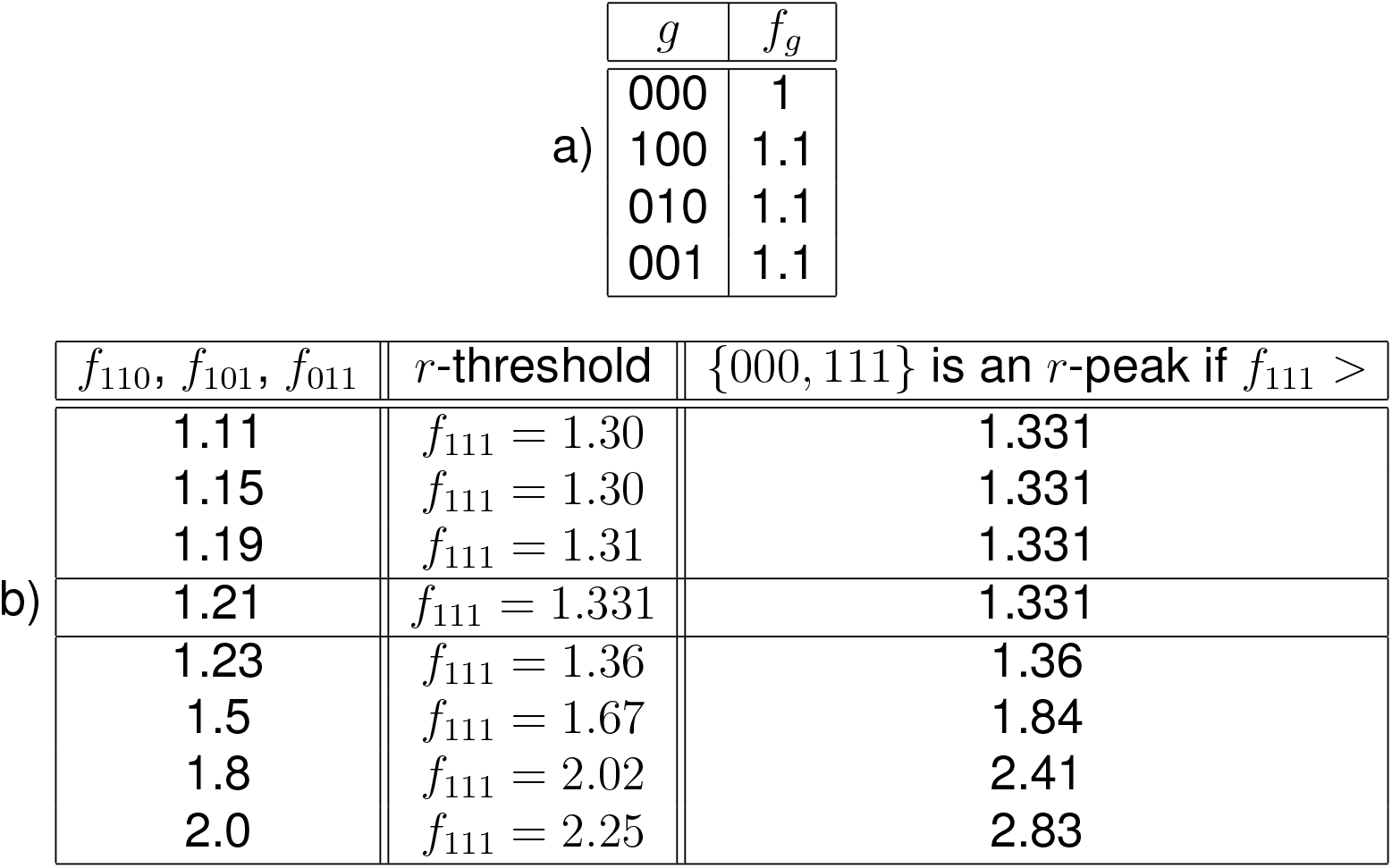
The fitness values are fixed for 000, 100, 010, 001 (Table 7a). The three double mutants have the same fitness value in each experiment (Table 7b). The tables show the recombination threshold along with the threshold that makes {000, 111} a recombination peak. Notice that fitness is multiplicative exactly if the double mutations have fitness 1.21.

|  |  |  |
| --- | --- | --- |
| a) | $g$ | $f_g$ |
|  | 000 | 1 |
|  | 100 | 1.1 |
|  | 010 | 1.1 |
|  | 001 | 1.1 |

|  |  |  |  |
| --- | --- | --- | --- |
| b) | $f_{110}, f_{101}, f_{011}$ | $r$-threshold | $\{000, 111\}$ is an $r$-peak if $f_{111} >$ |
| | 1.11 | $f_{111} = 1.30$ | 1.331 |
| | 1.15 | $f_{111} = 1.30$ | 1.331 |
| | 1.19 | $f_{111} = 1.31$ | 1.331 |
| | 1.21 | $f_{111} = 1.331$ | 1.331 |
| | 1.23 | $f_{111} = 1.36$ | 1.36 |
| | 1.5 | $f_{111} = 1.67$ | 1.84 |
| | 1.8 | $f_{111} = 2.02$ | 2.41 |
| | 2.0 | $f_{111} = 2.25$ | 2.83 |

**TABLE 8.** The set {000, 111} is a recombination peak if *f*_111_ > 1.65. The recombination threshold was also 1.65.

| $g$ | $f_g$ |
| --- | --- |
| 000 | 1 |
| 100 | 1.1 |
| 010 | 1.1 |
| 001 | 1.1 |
| 110 | 1.3 |
| 101 | 1.4 |
| 011 | 1.5 |
| $r$-threshold | $f_{111} = 1.65$ |

**TABLE 9.** The set {000, 111} is a recombination peak if *f*_111_ > 2.02. The recombination threshold was 1.88.

| $g$ | $f_g$ |
| --- | --- |
| 000 | 1 |
| 100 | 1.1 |
| 010 | 1.1 |
| 001 | 1.1 |
| 110 | 1.5 |
| 101 | 1.6 |
| 011 | 1.7 |
| $r$-threshold | $f_{111} = 1.88$ |

**TABLE 10.** The set {000, 111} is a recombination peak if *f*_111_ > 2.52. The landscape has substantial increasing returns epistasis except for the very last step.

| $g$ | $f_g$ |
| --- | --- |
| 0000 | 1 |
| 1000 | 1.1 |
| 0100 | 1.1 |
| 0010 | 1.1 |
| 0001 | 1.1 |
| 1100 | 1.3 |
| 1010 | 1.3 |
| 1001 | 1.3 |
| 0110 | 1.3 |
| 0101 | 1.3 |
| 0011 | 1.3 |
| 1110 | 2 |
| 1101 | 2 |
| 1011 | 2 |
| 0111 | 2 |
| $r$-threshold | $f_{111} = 2.23$ |

## Notes

### Competing Interest Statement

The authors have declared no competing interest.

### Summary of Updates

Some basic results on multi-allelic systems have been added in the beginning of the result section, as a service to the reader who may be new to the topic and need some overview. The new results are of independent interest. However, the main topic is still recombination.

